# Carbon limitation decouples roots but not leaves from nitrogen-fixing mutualists

**DOI:** 10.64898/2026.07.03.736439

**Authors:** Lucas José Reis Bartsch, Laura Carolina Leal, Anselmo Nogueira

## Abstract

While mutualistic symbioses with nitrogen-fixing bacteria enable plants to access fixed nitrogen, they also require substantial carbon investment. Under carbon limitation, such as shading, shifts in biomass allocation can decouple symbiotic investment from leaf and root growth, potentially compromising plant nitrogen status. Because shading shifts biomass allocation toward light acquisition, it could influence nitrogen fixing symbiosis in two opposing ways. If nodulation remains coupled to leaves rather than roots, nitrogen status should be maintained despite reduced root growth. Alternatively, if root growth constrains nodulation, nitrogen status should decline. We tested these hypotheses by manipulating light availability (full sunlight vs. 50% shade) and quantifying biomass allocation and symbiotic nodulation. Under shading, plants allocated proportionally more biomass to shoots than to roots and invested less biomass in root nodules. Relationships between nodulation and leaf or root biomass differed between treatments but converged with increasing plant size, although shaded plants never attained the root biomass observed in full sunlight. Leaf nitrogen concentration was maintained under shading because nodulation remained coupled to leaf investment despite reduced root allocation. These findings highlight that, under carbon limitation, maintaining leaf and nodule coupling enables plants to reduce nodule investment without compromising the nitrogen benefits of symbiosis.

**Highlight:** Under carbon limitation, maintaining leaf–nodule coupling despite reduced root investment enables plants to reduce nodule investment without compromising leaf nitrogen status.

## Introduction

Plants can establish mutualistic symbioses with nitrogen-fixing bacteria, thereby maintaining their performance in environments with low nitrogen availability (Zahran, 1999). In this interaction, nitrogen-fixing bacteria infect plant roots and induce the formation of root nodules, where they convert atmospheric nitrogen into ammonia, which becomes available to the host plant (Masson-Boivin *et al*., 2009; Terpolilli *et al*., 2012). Despite these benefits, symbiotic nodulation also imposes substantial carbon costson plants (Bronstein, 2001). Yet, these costs have received comparatively little attention in both intra- and interspecific studies (Bronstein, 2026). These costs result mainly from the continuous carbon inputs required to establish and maintain the partner bacteria (Schulze *et al*., 1999; Kiers and Denison, 2008; Lepetit and Brouquisse, 2023). When the carbon demand imposed by nodulation exceeds the plant’s carbon supply, the relative cost of the symbiosis increases and plant performance may decline (Voisin *et al*., 2013; Pringle, 2016a). Therefore, under reduced carbon availability, the balance between the costs and benefits of nitrogen-fixing symbiosis may shift, potentially reducing plant performance or even disrupting the mutualism.

Carbon limitation can alter how plants distribute resources among different plant functions, including the patterns of biomass allocation among plant organs and symbiotic root nodules. Under shaded conditions, for instance, plants commonly allocate proportionally more biomass to aboveground light-acquiring organs at the expense of root development (Reynolds and Thornley, 1982; Poorter *et al*., 2012). This can reduce the carbon available to support root symbionts and, consequently, its investment in nodulation (Chu and Robertson, 1974; Hansen *et al*., 1990; Xia, 1995; Araujo *et al*., 2018). In addition, the positive effects of the symbiosis on plant growth are often reduced under carbon limitation, even in nutrient-poor environments where the interaction with nitrogen-fixing bacteria tends to detrimental for plant (Lau *et al*., 2012; Friel and Friesen, 2019). However, variation in plant size can itself alter proportional patterns of carbon allocation among plant functions (Weiner, 2004; Liu *et al*., 2021). Consequently, apparent changes in allocation to nodules under carbon limitation may simply reflect differences in plant size rather than a reorganization of biomass allocation among plant organs (Weiner, 2004; Valladares *et al*., 2007). Disentangling these relationships is therefore essential to determine whether carbon limitation changes investment in symbiotic nodulation and the role of this symbiosis on plant’s performance.

The allocation patterns between nodulation and different plant organs may respond differently to carbon limitation. Because different organs impose distinct demands for carbon and nitrogen, nodulation may involve both allocation conflicts and/or functional coupling with other plant structures (Libault, 2014; Lepetit and Brouquisse, 2023). For example, leaves simultaneously represent the primary source of carbon and a major nitrogen sink in plants, with most of foliar nitrogen associated with photosynthetic proteins such as Rubisco (Gulmon and Chu, 1981; Mooney *et al*., 1981). Because carbon assimilation by leaves depends on nitrogen supply and nitrogen fixation in nodules depends on carbon supply, investment in nodulation tends to scale with investment in leaves throughout plant growth (Takahashi *et al*., 2018; Kobayashi *et al*., 2021). In this case, the maintenance of plant nitrogen supply, and consequently, investment into aboveground organs may depend on preserving this leaf-nodule relationship at carbon-limiting conditions (Lepetit and Brouquisse, 2023). However, if carbon limitation constrains belowground investment, the resulting reduction in root growth may limit the amount of nodulation that can be supported (Desbrosses and Stougaard, 2011; Bensmihen, 2015). Under this scenario, restrictions on root investment could constrain the capacity of plants to maintain this relationship between leaf investment and symbiotic nitrogen acquisition, compromising its investment in light-acquiring organs.

In this experimental study, we investigated how changes in biomass allocation under carbon limitation affect the carbon costs and nutritional benefits of symbiotic nodulation in plants. Using shading to impose carbon limitation, we first evaluated its effects on whole-plant biomass allocation patterns and overall nodulation. Next, we examined how nodulation investment relates to leaf versus root biomass under contrasting light conditions. We hypothesized that shading would favor plant investment in light-acquiring organs (mainly leaves), favoring aboveground structures at the expense of belowground biomass. If the relationship between leaf investment and nodulation is maintained under carbon limitation, nodulation would remain coupled to leaf production despite reduced root growth, preserving plant nitrogen status. Alternatively, if nodulation is driven by plant root structure, reduced root investment under carbon limitation would constrain plant allocation to nodules, weakening the association between leaves and nodules, and reducing plant nitrogen status. By testing these alternative hypotheses, we aimed to determine how carbon limitation alters plant allocation strategies among leaves, roots and nodules, and the consequence of theses shifts for plant nitrogen status and ultimately, the maintenance of plant-fixing bacteria symbiosis.

## Material and Methods

### Focal plant species

We used the woody legume *Chamaecrista nictitans* (L.) Moench as our focal plant species. This annual plant is widely distributed throughout Brazil, occurring mainly in *Cerrado* semi-arid areas, including environments with low soil nitrogen availability (Prochazka *et al*., 2024; Rando *et al*., 2026). Its root system establishes symbiotic associations with nitrogen-fixing bacteria, typically *Bradyrhizobium* species (Santos *et al*., 2017; Casaes *et al*., 2023; Souza *et al*., 2025). The nodules resulting from this interaction exhibit indeterminate growth and a high degree of compartmentalization, a structural feature associated with the efficiency and stability of the symbiosis (Casaes *et al*., 2023). Due to this high level of specialization, *C. nictitans* strongly depends on its association with *Bradyrhizobium* to meet its nitrogen demands, exhibiting severe limitations in growth and development in the absence of this mutualistic symbiosis (Souza *et al*., 2025).

### Experimental design

To test our hypotheses, we cultivated 40 individuals of *C. nictitans* under experimental garden conditions. Seeds were obtained from nine mother plants belonging to two natural populations: Parque Estadual do Juquery (Franco da Rocha, São Paulo, Brazil) and the São Bernardo do Campo *campus* of the Federal University of ABC (São Bernardo do Campo, São Paulo, Brazil), ensuring genetic variability among experimental units. All seeds were mechanically scarified before sowing in late October. We planted at least three seeds per pot in 40 eight-liter pots containing a homogeneous mixture of 70% sand and 30% commercial substrate. This soil provided mineral nutrients while containing minimal amounts of nitrogen (Table S1). After germination, seedlings were thinned to one individual per pot when one of the seedlings reached four fully expanded leaves.

Plant growth was monitored for 120 days under full sunlight and well-watered conditions. When plants reached this age, they were assigned to two light treatments: full sunlight (0% shading, n = 19) and 50% shading (n = 20), remaining under these conditions for approximately 40 days. One plant in the full-sunlight treatment died during the experiment and was excluded from the analyses, resulting in the final sample size of 19 plants. Before treatment application, we ensured that plants assigned to each treatment had similar sizes, avoiding any size bias between treatments (Table S2). To manipulate light availability, we constructed rectangular wooden frames using thin wooden slats. We used shade cloth reducing 50% of incident light in the shaded treatment, applied only to the top and upper lateral portions of the frames. For full-light treatment, the frames were completely uncovered. In the 50% shade treatment, we created openings along the lower lateral portions of the frames to allow air circulation and minimize differences in ventilation and temperature between treatments. Finally, we allocated five plants to each wood frame of each treatment (block). At least one individual from each of the nine maternal plants was allocated to each treatment.

Plants were distributed across two wooden tables, with shaded plants placed on one table and full-sun plants on the other to avoid interference of the shading structures on control plants. The tables were positioned so that both treatments received similar exposure to morning and late-afternoon sunlight. Individual plant positions within each table were randomized. The experiment was conducted during the summer, between February and March, under naturally warm conditions and frequent rainfall. Plants were manually watered twice daily with 300 mL of water per pot. Each plant was considered an experimental unit.

### Carbon allocation measurements

To investigate if carbon limitation induced by shading would shift biomass allocation toward light-acquisition structures at the expense of root growth, and whether nodulation scaled more closely with leaf or root investment under these conditions, we measured mean leaf area, the biomass of above- and belowground organs, and nodule traits under both light treatments. After 40 days under the treatments, plants were harvested, and leaves, stems, roots, and root nodules of each individual were manually separated. The number of root nodules was then recorded for each plant. We randomly chose and scanned three fully expanded leaves from each plant to obtain the mean leaf area per plant. Then, we dried all plant organs at 50 °C for ten days and weighed daily the samples using an analytical balance (0.0001 g precision) from the fifth day onward until constant mass was reached. We used the dry mass of leaves (including the three leaves used for leaf area estimation), stems, roots, and nodules as a proxy for carbon allocation to each structure, and the ratio between aboveground (leaves + stems) and belowground (roots + nodules) biomass as an indicator of whole-plant carbon partitioning.

### Nitrogen content measurements

To investigate whether shading influences the leaf nitrogen content of plants, we measured leaf nitrogen content under both light treatments. For each plant, we collected a pooled sample of 10 leaves, systematically selected from different branches and positions along the shoots of each plant to capture variation in leaf age. Leaves were first dried, and then the leaflets were detached, lyophilized and ground into a fine homogeneous powder. We weighted two subsamples (2.6–2.8 mg) of the powder material from each plant, encapsulated it in tin capsules, and analyzed it nitrogen content using an elemental analyzer (Flash EA 1112, Thermo Scientific, USA) to determine leaf nitrogen content. We expressed the nitrogen content as percentage of dry mass (%N) that was then converted in an mg g^-1^ concentration of nitrogen.

### Statistical Analysis

To test our hypothesis that carbon limitation induced by shading would shift biomass allocation toward light-acquisition structures at the expense of root growth, and to investigate whether nodulation scaled more closely with leaf or root investment under these conditions, we used standardized major axis regressions (SMA). This allometric approach is widely used to estimate scaling exponents (β, hereafter referred to as slopes) and allometric elevation constants (γ, hereafter referred to as intercepts) describing relationships among biological traits (Warton *et al*., 2006). For each relationship, we fitted the allometric equation Y = γX^β^, where X and Y represented the traits being compared. Four allometric relationships were evaluated: aboveground mass versus belowground mass, leaf dry mass versus total plant dry mass, nodule dry mass versus leaf dry mass, and nodule dry mass versus root dry mass. We also evaluated the relationship between mean leaf area and total plant dry mass because shaded plants may increase leaf area independently of changes in leaf and plant biomass. This complementary analysis allowed us to detect morphological adjustments that may enhance light interception of leaves under carbon limitation.

We included shading treatment as a categorical factor in all models described above to test whether slopes and intercepts differed between light conditions. We first tested for differences in slope. When slopes did not differ between full-light and shaded plants, we fitted a common slope model to test for differences in intercept. We used log_10_-transformed data to linearize the allometric relationships as Log_10_(Y) = Log_10_(γ) + βLog_10_(X). Using the linearized relationships, we tested for differences in slope and intercept between light treatments. Differences in slope indicate changes in how one trait scales with another, whereas differences in intercept indicate greater or lower investment in one trait relative to the other for a given value of the reference trait.

To determine whether shading affected plant investment in symbiotic nodulation and plant nitrogen content, we fitted three linear mixed-effects model (LMM) with shading treatment as the predictor variable and either nodule number, nodule dry mass and leaf nitrogen concentration as the response variable. Seed maternal origin and block (each wood frame with five plants each) were included as random effects. The significance of the fixed effect was assessed using Satterthwaite’s t-tests. If carbon limitation reduces investment in nodulation, we expected shaded plants to produce fewer nodules and lower total nodule mass than plants under full sunlight. Also, if nodulation continues to scale with leaf investment under carbon limitation, leaf nitrogen concentration should not differ between shaded and full sunlight plants. Alternatively, if reduced root growth constrains nodulation, shaded plants should exhibit lower leaf nitrogen concentration than plants grown under full sunlight.

We performed all analyses in R version 4.4.3. We used the *smatr* package (Warton *et al*., 2012) for standardized major axis analyses and the *lme4* (Bates *et al*., 2014) and *lmerTest* packages (Kuznetsova *et al*., 2017) for fitting the linear mixed-effects models.

## Results

Shading reduced in about 53% both nodule dry mass per plant (t = −4.788, df = 27.44, p < 0.001; Figure S1A, Table S3) and in about 57% the number of nodules per plant (t = −5.109, df = 29.28, p < 0.001; Figure S1B, Table S3). We observed no differences in slopes of allometric relationships between aboveground and belowground biomass (LR = 1.26, p = 0.26, Figure 1A, Table S4), leaf biomass and total biomass (LR = 0.24, p = 0.62, Figure 1B, Table S4) and leaf area and total biomass (LR = 0.54, p = 0.81, Figure 1C, Table S4) between full-light and shaded plants. However, shaded plants exhibited higher allometric intercept for aboveground versus root biomass (Wald = 17.27, p < 0.001, Figure 1A, Table S4) and leaf area versus total biomass (Wald = 33.3, p < 0.001, Figure 1C, Table S4), indicating greater plant investment in light-capturing tissues, as expected. A small difference in allometric intercept was also detected for leaf biomass versus total biomass (Wald = 6.67, p = 0.009, Figure 1B, Table S4), corresponding to an increase of only 8%.

**Figure 1.**
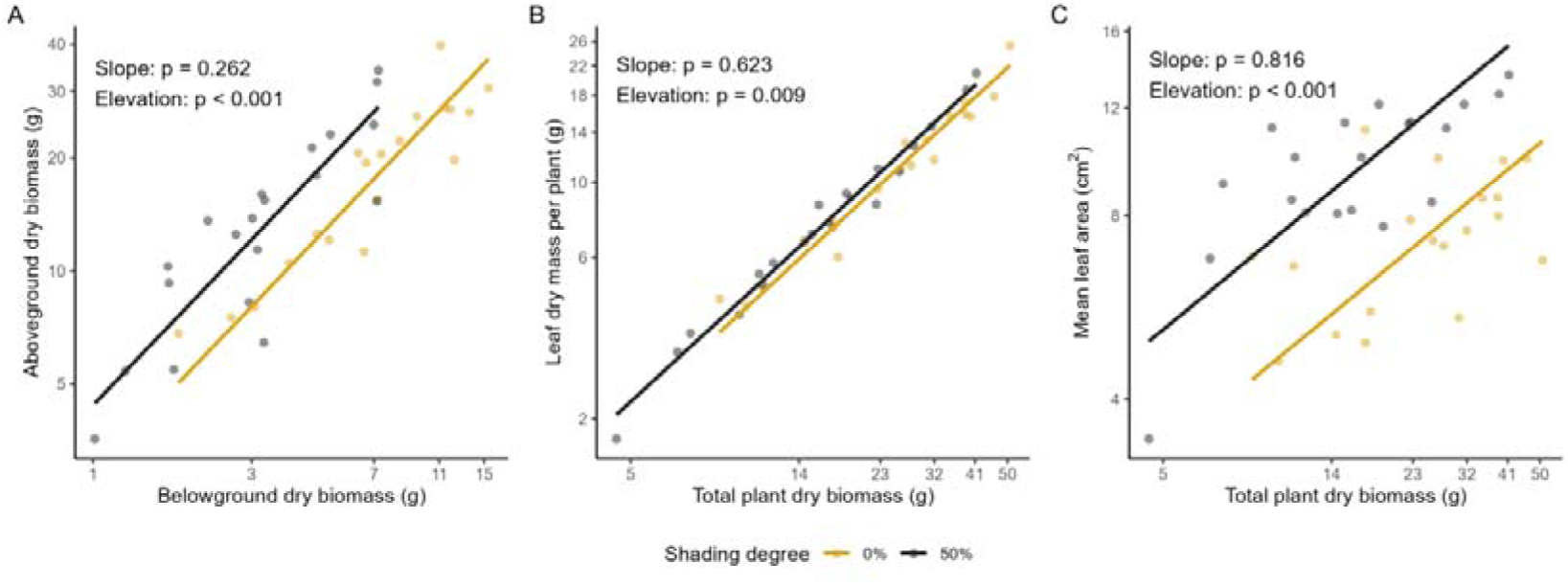
Relationships between belowground and aboveground dry biomass (A), total biomass and leaf area (B), and total biomass and leaf dry mass (C) in *Chamaecrista nictitans* (Fabaceae) grown under full sunlight (0%, yellow, n = 19) and 50% shading (gray, n = 20). All axes are log_10_-transformed. Points represent individual plants, and lines represent the fitted standardized major axis (SMA) regressions used to assess biomass allocation and leaf display responses to shading. Panels display the *P*-value for the likelihood ratio test comparing slopes between shading treatments and, when slopes did not differ significantly, the *P*-value for the Wald test comparing intercepts.

The relationship between nodule mass and leaf mass exhibited a higher allometric positive slope under shading (β = 1.41 ± 0.22, Figure 2A, Table S4) than under full sunlight (β = 0.94 ± 0.22; LR = 7.62, p < 0.005, Figure 2A, Table S4), indicating that shading increased the positive relationship between leaves and nodules. Likewise, the relationship between nodule and root mass exhibited a higher allometric intercept under shading (β = 1.50 ± 0.43, Figure 2B, Table S4) than under full sunlight (β = 0.79 ± 0.11; LR = 13.31, p < 0.001, Figure 2B, Table S4), indicating that nodules scaled more strongly with roots under carbon limitation. Leaf nitrogen concentration did not differ between shading treatments (t = 1.68, df = 5.78, p = 0.145, Figure 2C, Table S5).

**Figure 2.**
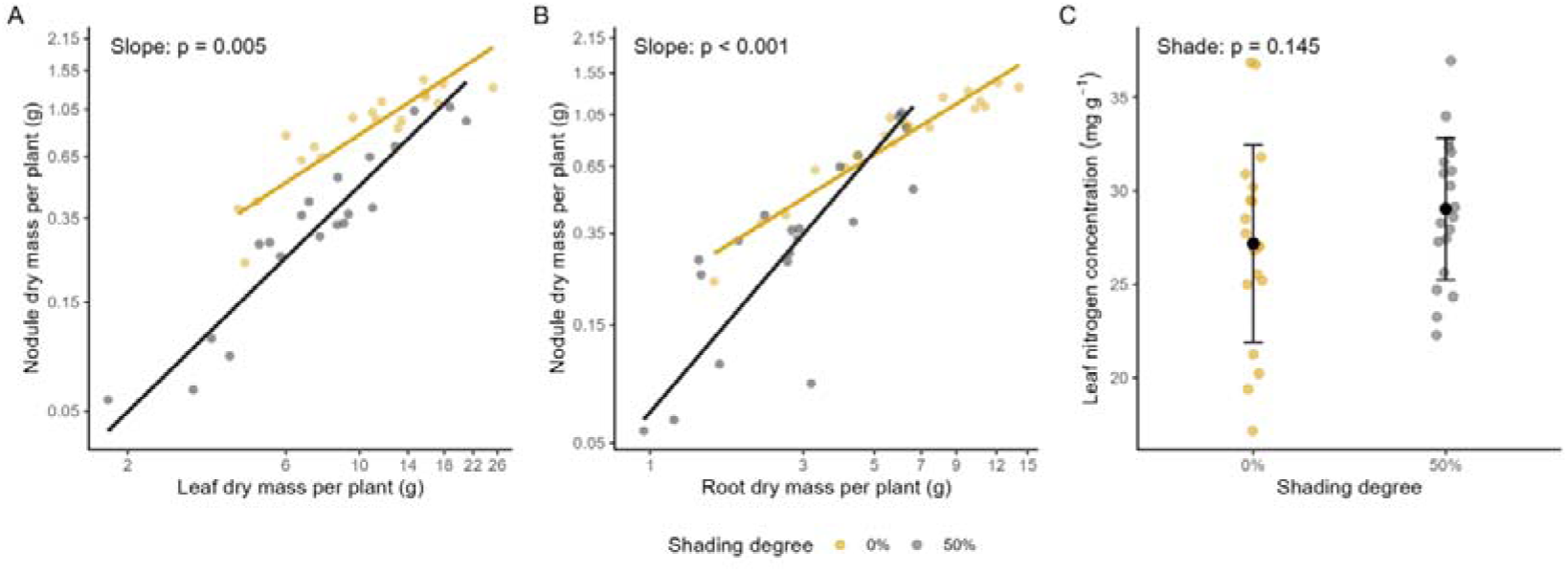
Effects of shading on the symbiosis between *Chamaecrista nictitans* (Fabaceae) and nitrogen-fixing bacteria. Relationships between nodule dry mass and leaf dry mass (A) and root dry mass (B) in plants grown under full sunlight (0%, yellow, n = 19) and 50% shading (gray, n = 20). Points represent individual plants and lines represent fitted standardized major axis (SMA) regressions. All axes are log[[-transformed. Panels display the *P*-value for the likelihood ratio test comparing slopes between shading treatments and, when slopes did not differ significantly, the *P*-value for the Wald test comparing intercepts. (C) Leaf nitrogen concentration in plants grown under full sunlight (0%, yellow) and 50% shading (gray). Points represent individual plants, black circles indicate treatment means, and error bars represent ± 1 SD. Asterisks indicate significant differences between treatments based on Satterthwaite’s *t*-test.

## Discussion

Symbiotic associations with nitrogen-fixing bacteria provide plants with access to biologically fixed nitrogen (Zahran, 1999) but require substantial carbon investment by the host plant (Pringle, 2016a). Here, we hypothesized that plants under shade would prioritize light acquisition over root growth, with symbiotic nodulation remaining coupled to leaf investment (Hypothesis 1, preserving nitrogen status) or to root investment (Hypothesis 2, reducing nitrogen status). Our results partially support Hypothesis 1, as we observed a stronger positive relationship between nodule biomass and leaf biomass than with root biomass in shaded plants, indicating that carbon allocation to symbiotic nodulation was more closely associated with investment in carbon-acquiring tissues rather than belowground biomass. Despite these shifts in allometric scaling under carbon limitation, leaf nitrogen concentration remained unchanged, demonstrating that nitrogen acquisition was maintained. Although shaded plants never attained the largest root systems observed under full sunlight, they achieved comparable maximum levels of leaf and nodule mass, suggesting that maintaining investment in symbiotic nodulation is a key plant strategy for sustaining nitrogen acquisition under carbon limitation. Together, these results indicate that (1) carbon limitation reshaped the allometric relationships among leaves, roots, and nodules; (2) under shading, nodule biomass scaled more closely with leaf biomass than with root biomass; and (3) these allocation shifts preserved plant nitrogen status despite reduced carbon availability.

In shaded plants, lower leaf investment was linked to disproportionately lower nodule investment, revealing that the leaf–nodule relationship was not consistent across the full range of leaf biomass. This pattern is consistent with the mathematical model proposed by Kobayashi et al. (2021), which predicts reduced nodule production per unit of leaf biomass under low-light conditions. Given the high carbon costs of biological nitrogen fixation, shaded plants with low leaf investment may be unable to sustain the same level of nodulation observed in plants grown under full sunlight (Schwember *et al*., 2019; Kobayashi *et al*., 2021). In contrast, for a given investment in leaf biomass, shaded plants developed proportionally greater leaf area. This is a common plant response to low light availability because it enhances light interception and increases the efficiency of leaf biomass investment for carbon acquisition (Evans and Poorter, 2001; Poorter *et al*., 2009). Consequently, although leaf biomass remained similar between treatments, shaded plants likely achieved greater carbon assimilation capacity per unit of leaf biomass, as greater leaf area increases whole-plant photosynthetic capacity (Peterson and Zelitch, 1982; Koyama and Kikuzawa, 2009). This increase may have enhanced their capacity to support the high carbon costs of biological nitrogen fixation (Sheehy and Phillips, 1987; Pringle, 2016a), allowing nodule investment to increase rapidly once sufficient leaf biomass was attained. Consequently, although plants with low leaf investment exhibited strong constraints on nodulation, those with greater leaf investment reached levels of nodule investment comparable to those of full-light plants for a given investment in leaf biomass.

Although plant investment in nodules followed investment in aboveground organs more closely than investment in belowground organs, nodule biomass responded more strongly to root biomass in shaded plants than in plants grown under full sunlight. Studies investigating the relationship between root investment and nodulation under carbon limitation caused by shading have reported either the maintenance of a proportional relationship (Chu and Robertson, 1974; Houx *et al*., 2009) or a reduction in the proportion of nodules per unit of root biomass (Friel and Friesen, 2019). Our results suggest that these contrasting findings may reflect the influence of plant size on this relationship (Weiner, 2004). In smaller shaded plants, nodulation was disproportionately low relative to root investment. However, this difference decreased as plant size increased, so that, at the highest levels of root investment under shading, the relationship between roots and nodules approached the same levels observed in plants grown under full sunlight. Importantly, this convergence occurred even though shaded plants never attained the highest root investment levels observed in full sunlight. This pattern indicates that limited root investment did not prevent larger shaded plants from establishing root–nodule relationships comparable to those of plants grown under full sunlight. In light of these findings, although several studies have shown that shading simultaneously reduces investment in roots and nodules (Chu and Robertson, 1974; Houx *et al*., 2009; Lau *et al*., 2012; Friel and Friesen, 2019), our results provide evidence that this association reflects carbon limitation related to reduced whole-plant growth, rather than reduced root investment per se. Together with the stronger dependence of nodulation on leaf investment, this biomass allocation shifts highlight that carbon availability exerts a stronger influence on nodule investment than root development under carbon limitation.

Despite the stronger dependence of nodule investment on leaf investment under shading, shaded plants still produced fewer nodules per unit of leaf biomass than plants under full sunlight. Nevertheless, leaf nitrogen concentration was maintained at similar levels in shaded plants and those grown under full sunlight. This apparent mismatch between reduced nodule investment and maintained nitrogen content aligns with evidence that nitrogen fixation efficiency per unit nodule biomass can be maintained (Chu and Robertson, 1974; Houx *et al*., 2009) or even increased (Santos *et al*., 1997) under shading, reinforcing that reduced nodule mass investment does not necessarily prevent nitrogen fixation from remaining sufficiently efficient to meet plant nitrogen requirements. The adjustment between leaf and nodule investment allows nodulation to be reduced to an extent that does not compromise the plants’ nitrogen status (Voisin *et al*., 2013; Kobayashi *et al*., 2021). Together, these findings indicate that plants can reduce the carbon costs of nitrogen-fixing symbiosis while maintaining much of its nutritional benefit under carbon limitation.

## Conclusion

Plants can reduce the carbon costs of nitrogen-fixing symbiosis without proportionally reducing its nutritional benefits under carbon limitation. This decoupling between investment and functional outcome highlights how resource limitation reshapes the cost–benefit balance of mutualistic interactions. Plants maintained the relationship between leaf and nodule investment despite reduced investment in roots, and reduced nodulation did not compromise leaf nitrogen status, indicating that the symbiosis remains functional even under carbon restriction. We show here that, although carbon availability is a major factor driving the cost–benefit balance of plant mutualistic interactions as a whole (Pringle, 2016a,*b*), the outcome of these interactions remains positive, mainly due to changes in patterns of plant carbon acquisition and allocation to symbiotic partners. It contributes to a growing body of evidence showing that mutualisms are shaped by resource limitation through its effects on allocation strategies and on the balance between costs and benefits. It speaks directly to other studies demonstrating that mutualisms generally remain beneficial across ecological contexts, only rarely shifting to parasitism or becoming destabilized by conflicts between partners (Frederickson, 2017; Leal *et al*., 2026). Our findings suggest that maintaining the relationship between leaf and nodule investment may represent a mechanism by which plant–rhizobium symbiosis remains beneficial under carbon limitation. Moreover, they support the view that mutualism robustness stems from the ability of partners to adjust resource allocation under environmental change without compromising the benefits of the interaction. This provides new insight into the mechanisms that sustain mutualisms in heterogeneous environments, and highlights the importance of integrating resource allocation and symbiotic costs into predictive frameworks for mutualism functioning under environmental change.

## Supplementary data

Figure S1. Effects of shading on nodule dry mass per plant (A) and the number of nodules per plant (B).

Table S1. Chemical and physical properties of the soil used in the experiment, including nutrient availability, soil acidity, cation exchange capacity, micronutrient concentrations, particle-size distribution, and total nitrogen content.

Table S2. Results of linear mixed-effects models testing whether plants assigned to different shading treatments differed in initial size before the experiment.

Table S3. Results of linear mixed-effects models testing the effects of shading on nodule dry mass and the number of nodules per plant.

Table S4. Results of standardized major axis analyses evaluating the effects of shading on allometric relationships among plant growth traits and plant–rhizobium symbiosis traits.

Table S5. Results of a linear mixed-effects model testing the effect of shading on leaf nitrogen concentration.

## Acknowledgements

We would like to thank Taiguara Ortiz and Luana Prochazka for providing the location and assisting in the final processing of the experiment’s data and express our gratitude to Samantha Marcon for her support in the field experiment and the schematic of the hypothesis in the Figure 1. We acknowledge the support of the Multiuser Center for Biodiversity and Conservation (CMBC-PROPES) at UFABC, which provided equipment and technical infrastructure for conducting the experiment. The authors also would like to disclosure the use of IA (Deepseek) for checking English grammar issues and improve the flow of some specific sentences along the manuscript.

## Author Contributions

L.J.R.B., and A.N conceived the ideas and designed methodology; L.J.R.B. collected the data; L.J.R.B. analysed the data; L.J.R.B., L.C.L. and A.N. led the writing of the manuscript. All authors contributed critically to the drafts and gave final approval for publication.

## Data Availability

Data and R scripts supporting the findings of this study are available in a private repository for peer review at: https://figshare.com/s/b99b57613cea185a6bee. The repository includes raw data tables, processed datasets, metadata, and all scripts used for statistical analyses and figure generation. Upon acceptance, all materials will be openly available in Figshare at 10.6084/m9.figshare.32897084.

---

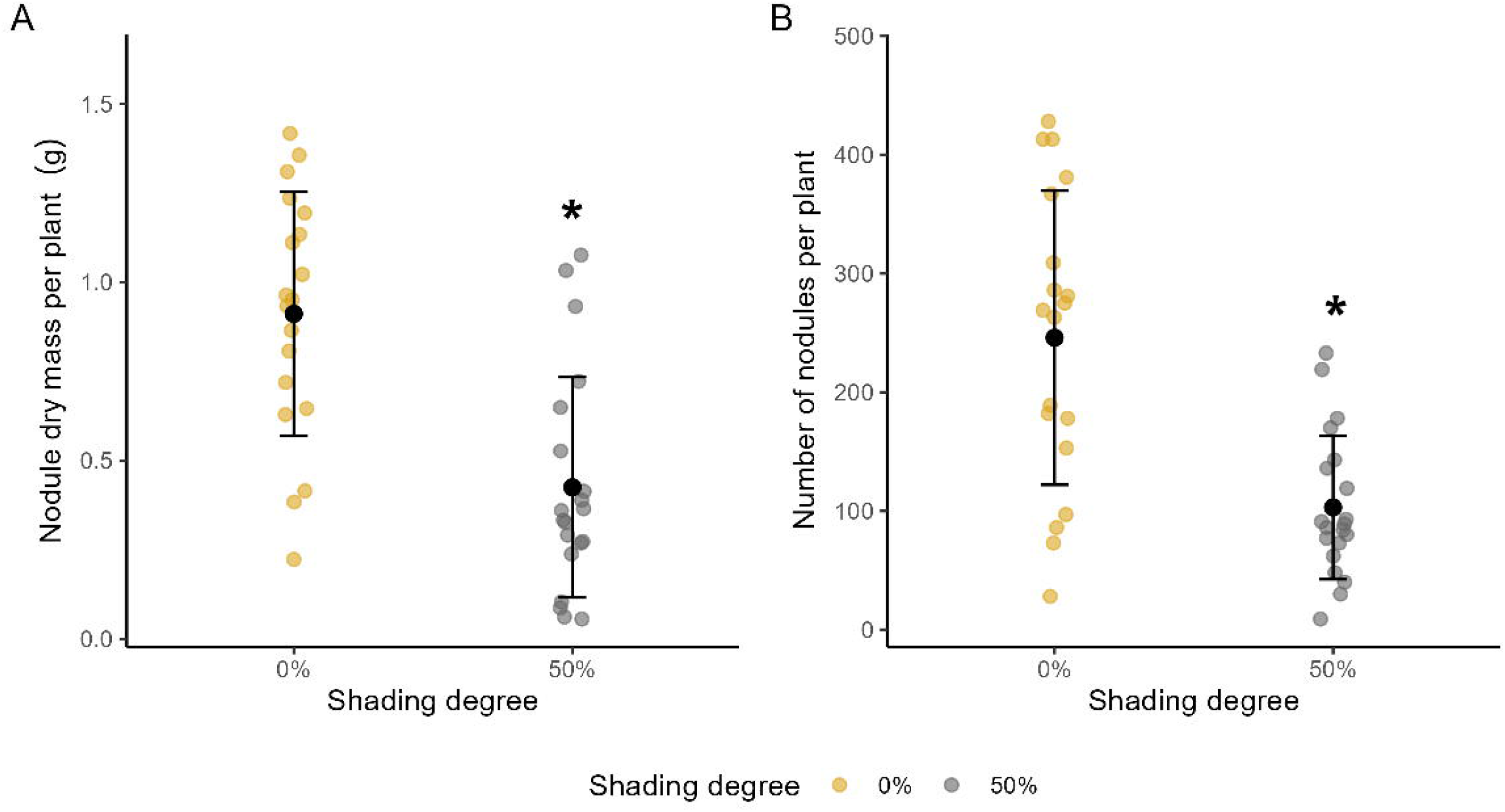

## Figure Alt Texts

Figure 1. Three-panel figure comparing allometric relationships between plants grown under full sunlight (yellow) and 50% shading (gray). Panel A shows aboveground dry biomass versus belowground dry biomass. Panel B shows leaf dry mass versus total plant dry biomass. Panel C shows mean leaf area versus total plant dry biomass. Each panel displays individual plants and fitted regression lines for each shading treatment. Slopes do not differ significantly between treatments in any panel. Shaded plants show higher intercept in panels A and C, indicating greater aboveground biomass and mean leaf area for a given plant size. In panel B, the fitted lines are nearly overlapping, with only a slight but statistically significant elevation difference between treatments.

Figure 2. Three-panel figure comparing plants grown under full sunlight (yellow) and 50% shading (gray). Panels A and B are scatterplots with fitted regression lines showing positive relationships between nodule dry mass and leaf dry mass (A) and between nodule dry mass and root dry mass (B). In both panels, the regression slopes differ significantly between shading treatments, with shaded plants showing steeper relationships. Panel C is a dot plot of leaf nitrogen concentration for the two shading treatments, with black circles indicating treatment means and error bars representing standard deviations. Mean leaf nitrogen concentration is similar between treatments, and no significant difference is detected.

